# Carbohydrate-Binding Protein from Stinging Nettle as Fusion Inhibitor for SARS-CoV-2 Variants of Concern

**DOI:** 10.1101/2022.07.08.499297

**Authors:** Emiel Vanhulle, Thomas D’huys, Becky Provinciael, Joren Stroobants, Anita Camps, Sam Noppen, Dominique Schols, Els J.M. Van Damme, Piet Maes, Annelies Stevaert, Kurt Vermeire

## Abstract

*Urtica dioica* agglutinin (UDA) is a carbohydrate-binding small monomeric protein isolated from stinging nettle rhizomes. It inhibits replication of a broad range of viruses, including coronaviruses, in multiple cell types, with appealing selectivity. In this work, we investigated the potential of UDA as a broad-spectrum antiviral agent against SARS-CoV-2. UDA potently blocks entry of pseudotyped SARS-CoV-2 in A549.ACE2^+^-TMPRSS2 cells, with IC_50_ values ranging from 0.32 to 1.22 µM. Furthermore, UDA prevents viral replication of the early Wuhan-Hu-1 strain in Vero E6 cells (IC_50_ = 225 nM), but also the replication of SARS-CoV-2 variants of concern, including Alpha, Beta and Gamma (IC_50_ ranging from 115 to 171 nM). In addition, UDA exerts antiviral activity against the latest circulating Delta and Omicron variant in U87.ACE2^+^ cells (IC_50_ values are 1.6 and 0.9 µM, respectively). Importantly, when tested in Air-Liquid Interface (ALI) primary lung epithelial cell cultures, UDA preserves antiviral activity against SARS-CoV-2 (20A.EU2 variant) in the nanomolar range. Surface plasmon resonance (SPR) studies demonstrated a concentration-dependent binding of UDA to the viral spike protein of SARS-CoV-2, suggesting interference of UDA with cell attachment or subsequent virus entry. Moreover, in additional mechanistic studies with cell-cell fusion assays, UDA inhibited SARS-CoV-2 spike protein-mediated membrane fusion. Finally, pseudotyped SARS-CoV-2 mutants with N-glycosylation deletions in the S2 subunit of the spike protein remained sensitive to the antiviral activity of UDA. In conclusion, our data establish UDA as a potent and broad-spectrum fusion inhibitor for SARS-CoV-2.

## 1 Introduction

The severe acute respiratory syndrome coronavirus 2 (SARS-CoV-2) swiftly spread from the 41 initially reported patients in Hubei province, China (1), to a global pandemic with 549 million confirmed cases (https://covid19.who.int/) and an estimated 18.2 million excess deaths in two years (2). Undoubtedly, Coronavirus Disease-2019 (COVID-19) presents an immense threat to the public health worldwide and the global economy. Several vaccines have already been approved, but worldwide vaccination coverage is still insufficient. In addition, current vaccines are suboptimal in preventing transmission, and novel variants of the virus with reduced susceptibility to the vaccines continue to emerge (3). Antivirals are a critical addition to the vaccination campaigns, to increase the resilience to SARS-CoV-2 infection, particularly in at-risk populations. Currently authorised COVID-19 therapeutics include remdesivir (4), ritonavir-boosted nirmatrelvir (Paxlovid) (5), molnupiravir (6), and certain anti-SARS-CoV-2 monoclonal antibodies (7).

SARS-CoV-2 entry in the host cell is mediated by its spike (S) protein, which is post-translationally cleaved into two subunits. The receptor-binding domain (RBD), which recognizes the angiotensin-converting enzyme 2 (ACE2) receptor (8), is located in the S1 subunit, while the S2 subunit harbours the fusion machinery. Cleavage at the S2′ site, which renders the spike protein fusion-competent, occurs for most SARS-CoV-2 variants preferably at the cell surface by type II transmembrane serine proteases (TTSP) such as transmembrane protease serine 2 (TMPRSS2) (9). In contrast, the Omicron variant favours the alternative endosomal entry pathway, where fusion activation depends on cathepsins (10, 11). About 40 % of the spike protein surface is decorated by glycans, which shield the virus from the host innate immune system. In total, 22 *N*-linked glycosites and 17 *O*-glycosites were identified on the SARS-CoV-2 spike (12). Glycans mediate protein folding and facilitate immune evasion (13), impact viral infectivity (14, 15), spike stability, and processing of the viral envelope protein by host proteases (16, 17). Glycans can be bound by mammalian lectins (carbohydrate-binding proteins) which are often expressed on immune and endothelial cells and are involved in virus internalization and transmission (18). In addition, many non-mammalian lectins are endowed with antiviral activity.

Previous results from our research group established plant lectins as a unique class of antiviral molecules (18). One of those, *Urtica dioica* agglutinin (UDA), a lectin isolated from stinging nettle rhizomes, is a small (8.5 kDa) monomeric protein with high glycine, cysteine and tryptophan content (19-21). It comprises two hevein-like domains, each with a saccharide-binding site (22), and exhibits carbohydrate-binding specificity for N-acetylglucosamine oligomers as well as high-mannose-type N-glycans (23, 24). UDA displays low cytotoxicity and potent antiviral activity against a wide spectrum of viruses; some examples are: human immunodeficiency virus (HIV) (25), cytomegalovirus (CMV) (25), respiratory syncytial virus (RSV) (25), influenza A and B virus (23, 25, 26), hepatitis C virus (27), herpes simplex virus (HSV) (23), dengue virus (DENV) (23, 28), and severe acute respiratory syndrome coronavirus (SARS-CoV) (29). Previous experiments with HIV, RSV and influenza virus have shown that UDA interferes with virus entry, presumably by hindering virus fusion (25, 26).

The ongoing COVID-19 pandemic highlights the need for a broad-spectrum antiviral that can be used immediately to rapidly diminish viral spread when an epidemic with a (re-)emerging virus occurs. We aimed to further explore and understand the broad and potent antiviral activity of UDA. Candidate SARS-CoV-2 entry inhibitors should be able to target both endosomal and TTSP-mediated cell surface entry, and show broad activity against different variants in different cell types. Hence, resolving its precise mechanism of action in detail is pivotal to decide whether the lectin is a potential broad-acting antiviral inhibitor. In the present study, we evaluate UDA against a panel of SARS-CoV-2 variants in different cell types and demonstrate a consistent antiviral activity of UDA against SARS-CoV-2. We propose membrane fusion as the possible target for antiviral intervention, identifying UDA as entry/fusion inhibitor for SARS-CoV-2.

## 2 Materials and methods

### 2.1 Cell lines, primary cells and virus strains

#### Cell lines

Human Embryonic Kidney 293T (HEK293T) cells (cat n° CRL-3216), African green monkey kidney Vero E6 cells (cat n° CRL-1586) and human adenocarcinomic alveolar epithelial cells A549 (cat n°CCL-185) were obtained from ATCC (Manassas, VA, USA) as mycoplasma-free stocks. Together with an in-house designed human glioblastoma cell line, stably expressing ACE2 (U87.ACE2^+^) (30), these cells were grown in Dulbecco’s Modified Eagle Medium (DMEM, Thermo Fisher Scientific (TFS)) supplemented with 10% (*v/v*) fetal bovine serum (FBS; Hyclone). A549 lung carcinoma cells expressing human ACE2 and human TMPRSS2 (A549.ACE2^+^.TMPRSS2^+^; cat n° a549-hace2tpsa, Invivogen) were grown in DMEM/10% FBS supplemented with 100 µg/ml Normocin, 0.5 µg/ml Puromycin and 300 µg/ml Hygromycin. Cell lines were maintained at 37°C in a humidified environment with 5% CO_2_ and passaged every 3-4 days.

#### Primary cells and air-liquid interface cultures

SmallAir™ (cat n° EP21SA) and MucilAir™ (cat n° EP01MD, bronchial cell origin) were purchased from Epithelix Sàrl (Geneva, Switzerland) and maintained in SmallAir™ medium (cat n° EP65SA) and MucilAir™ medium (cat n° EP05MM), respectively. Medium of the ALI cultures was changed every other day and transepithelial/transendothelial electrical resistance (TEER) was measured on a regular base.

#### Viruses

All virus-related work was conducted in the high-containment biosafety level 3 facilities of the Rega Institute from the Katholieke Universiteit (KU) Leuven (Leuven, Belgium), in accordance with institutional guidelines. Severe acute respiratory syndrome coronavirus 2 (SARS-CoV-2) isolates were recovered from nasopharyngeal swabs of RT-qPCR-confirmed human cases obtained from the University Hospital (Leuven, Belgium). SARS-CoV-2 viral stocks were prepared by inoculation of confluent Vero E6 cells in DMEM supplemented with 2% (*v/v*) FBS, as described in detail (31). Recombinant SARS-CoV-2-GFP virus (Wuhan strain), as described in (32), was a kind gift of Dr. Volker Thiel (University of Bern, Switzerland). Titers were determined by tissue culture infectious dose 50 (TCID_50_) method of Reed and Muench (33) on Vero E6 and U87.ACE2^+^ cells. Viral genome sequence was verified, and all infections were performed with passage 3 to 5 virus.

### 2.2 Antibodies and compounds

#### Antibodies

The following antibodies were used for western blotting: ACE2 Polyclonal Goat IgG (cat. n° AF933, R&D systems), anti-β-actin (cat. n° MA1-140, Invitrogen), HRP-labelled goat anti-mouse immunoglobulin (IgG; cat. n° P0447, Dako). The following antibodies were used for flow cytometry: rabbit polyclonal SARS-CoV-2 nucleocapsid-specific antibody (cat. n° GTX135357, GeneTex), rabbit monoclonal SARS-CoV-2 spike-specific antibody [R001] (cat. n° 40592-R001, Sino Biological), mouse monoclonal SARS-CoV-2 spike-specific antibody [MM57] (cat. n° 40592-MM57, Sino Biological), mouse monoclonal SARS-CoV/SARS-CoV-2 spike-specific antibody [1A9] (cat. n° GTX632604, GeneTex), Alexa Fluor 647 (AF647)-labelled goat anti-rabbit IgG polyclonal antibody (cat. n° 4414, Cell Signaling Technologies), and phycoerythrin (PE)-labelled goat anti-mouse IgG (cat. n° 405307, BioLegend). The following antibodies were used for surface plasmon resonance studies: purified SARS-CoV-2 spike-specific monoclonal rabbit primary antibodies R001 and R007 (cat. n° 40592-R001 and cat. n° 401_50_-R007, Sino Biological)

#### Compounds

*Urtica dioica* agglutinin (UDA) from Stinging Nettle was from EY Laboratories, CA, USA (cat. n° L-8005-1).

### 2.3 Plasmid construction

pCAGGS.SARS-CoV-2_SΔ19_fpl_mNG2(11)_opt was generated using NEBuilder DNA assembly (New England Biolabs) of a pCAGGS vector backbone cleaved using EcoRV-HF and HindIII-HF (New England Biolabs) and a PCR fragment encoding a codon-optimized SARS-CoV-2 Wuhan-Hu-1 spike protein (amplified from pCMV3-C-Myc; VG40589-CM, SinoBiological) with a C-terminal 19 amino acid deletion as described in (34). A 12 amino acid flexible protein linker (fpl) and a modified 11^th^ betasheet of mNeonGreen (35) were added at the C-terminus. pCAGGS.BSD_fpl_mNG2(11) was generated using NEBuilder DNA assembly of a pCAGGS vector backbone cleaved using EcoRV-HF and HindIII-HF and the Blasticidin S deaminase gene (BSD) PCR amplified from a pLenti6.3 vector. Afterwards, cDNA encoding for a 12-amino acid fpl and a modified 11^th^ betasheet of mNeonGreen were inserted at the 3’
send of the insert ORF. pcDNA3.1.mNG2(1-10) was generated through NEBuilder DNA assembly of a pcDNA3.1 vector (TFS), amplified by PCR, and 10 betasheets of a modified mNeonGreen synthesized by Genscript. For pCAG3.1/SARS2-Sd19 PCR-amplified Wuhan-Hu-1 spike sequence (from pCMV3-C-Myc) was inserted via blunt end cloning in the pCAG3.1 acceptor vector cut with EcoRV-HF.

### 2.4 Immunoblotting

Immunoblotting analysis was performed as previously reported (30). Cells were collected and lysed in ice-cold NP-40 lysis buffer (50 mM Tris-HCL (pH 8.0), 150 mM NaCl, and 1% Nonidet P-40) supplemented with cOmplete Protease Inhibitor (Roche) and PMSF Protease Inhibitor (100 mM in dry isopropanol, TFS). Cell lysates were centrifuged at 17,000 *g* for 10 min at 4°C to pellet nuclei and debris. For SDS gel electrophoresis, supernatant samples were boiled in reducing 2x Laemmli sample buffer (120 mM Tris-HCl (pH 6.8), 4% SDS, 20% glycerol, 100 mM dithiothreitol, and 0.02% bromophenol blue). Equal volumes of lysate were run on Criterion XT Bis-Tris gels (4–12%; Bio-Rad) at 170 V for 55 min using 1x XT-MES buffer (Bio-Rad), transferred to nitrocellulose membranes using the BioRad Trans-Blot Turbo transfer system (Bio-Rad). Membranes were blocked for 1 h with 5% non-fat dried milk in TBS-T (20 mM Tris-HCL (pH 7.6), 137 mM NaCl, and 0.05% Tween-20). After overnight incubation with primary antibody at 4°C, membranes were washed and incubated for 1h with secondary antibody. β-actin was used as a loading control. SuperSignal West Pico and Femto chemiluminescence reagent (Thermo Fisher scientific) was used for detection with a ChemiDoc MP system (Bio-Rad). Signal intensities were quantified with Image Lab software v5.0 (Bio-Rad).

### 2.5 Wild type virus infection and antiviral assays

One day prior to the experiment, Vero E6, A549.ACE2^+^-TMPRSS2 and U87.ACE2^+^ cells were seeded in 96-well microtiter plates. Next day, 3- or 5-fold serial dilutions of the test compounds were prepared in virus infection media (same as cell culture medium, but with 2% FBS), overlaid on cells, and virus was added to each well (MOI indicated in the figure legends). Cells were incubated at 37°C under 5% CO_2_ for the duration of the experiment. At various timepoints p.i., the virus-induced cytopathic effect (CPE) and GFP expression was microscopically evaluated and GFP^+^ area was calculated as a percentage of the total cell area. Inhibition was calculated by comparison to virus control wells with no inhibitor added. IC_50_ values were determined by interpolation. In case of subsequent analysis to quantify viral genome copy numbers with RT-qPCR, infected cells were washed with PBS at 2h post-infection to remove unbound virus, followed by incubation with freshly prepared 3- or 5-fold serial dilutions of compounds (for antiviral assay) at 37°C, 5% CO_2_. At various timepoints, supernatants were collected and stored at -80°C until further analysis.

Four days after infection, the cell viability of mock- and virus-infected U87.ACE2^+^ cells was assessed spectrophotometrically via the in situ reduction of 3-(4,5-dimethylthiazol-2-yl)-5-(3-carboxy-methoxyphenyl)-2-(4-sulfophenyl)-2H-tetrazolium inner salt, using the CellTiter 96 AQueous One Solution Cell Proliferation Assay (Promega), as described before (30). The absorbances were read in an eight-channel computer-controlled photometer (Multiscan Ascent Reader, Labsystem, Helsinki, Finland) at two wavelengths (490 and 700 nm). The optical density (OD) of the samples was compared with sufficient cell control replicates (cells without virus and drugs) and virus control wells (cells with virus but without drugs). The concentration that inhibited SARS-CoV-2-induced cell death by 50% (IC_50_) was calculated from interpolation.

### 2.6 Virus infection of primary ALI cell cultures

Prior to infection, duplicates of SmallAir™ and MucilAir™ reconstituted bronchial epithelium were washed twice with PBS warmed to 37°C to remove mucus and debris and basal media were replenished with warm cell culture media. Compound was added simultaneously with 2 × 10^4^ TCID_50_ of SARS-CoV-2 20.EU2 strain or a GFP-encoding Wuhan-Hu-1 variant (theoretical MOI of 0.3) to the apical compartment. Compound and virus were diluted in DMEM supplemented with 2% FBS and incubated at 37°C, 5% CO_2_ for 2 h. Mock controls were exposed to the same volume of medium only. Subsequently, virus inoculum (with or without compound added) was removed and the apical compartment was washed twice with PBS to remove remaining unbound virus. The apical side of the ALI cultures were exposed to air till the end of the experiment, with an apical wash (with 200 µl PBS for 5 min at 37°C) at 24h p.i. Virus release was assessed in the apical wash at 4 days p.i.

### 2.7 Viral RNA extraction and reverse transcription quantitative PCR (RT-qPCR)

Supernatants and apical washes were harvested, viral particles were lysed and total RNA was extracted using QIAamp viral RNA mini kit (Qiagen, Switzerland) following manufacturer’s instruction. Viral RNA was quantified using a duplex RT-qPCR assay, using the QuantStudio™5 Real-Time PCR system (Applied Biosystems), which has been described in detail (31). Briefly, all primers and probes were obtained from Integrated DNA Technologies (IDT, Leuven, Belgium). Final concentration of combined primer/probe mix consist of 500 nM forward and reverse primer and 250 nM probe. Viral E and N genes are simultaneously amplified and tested using a multiplex RT-qPCR. All the procedures follow the manufacturer’s instructions of the Applied Biosystems TaqMan Fast Virus one-step mastermix (TFS). qPCR plate was read in the FAM and HEX channels using the following cycling protocol: 50°C for 5 min, 95°C for 20 sec, followed by 45 cycles of 95°C for 3 sec and 55°C for 30 sec. A stabilized *in vitro* transcribed universal synthetic single stranded RNA of 880 nucleotides in buffer with known copy number concentration (Joint Research Centre, European Commission, cat. n° EURM-019) was used as a standard to quantitatively measure viral copy numbers.

### 2.8 Immunofluorescence microscopy

U87.ACE2^+^ cells and ALI cultures of SmallAir™ cells were infected with a GFP-encoding SARS-CoV-2 variant. At indicated time-points, U87.ACE2^+^ cells and ALI cultures were imaged with a Primovert iLED inverted immunofluorescence microscope employing a 4X Plan-Achromat objective (Zeiss NTS Ltd). Representative images were captured in the green channel of the microscope to determine GFP expression. Images were processed and analyzed using the open-source image analysis software Fiji (36). In brief, images were added to a stack and converted to 8-bit. A threshold was set to separate background from GFP positive signal. GFP^+^ area was calculated as a percentage of the total cell area.

### 2.9 Cell-cell fusion assay

HEK293T and A549.ACE2^+^ cells were plated in 6-well plates to reach 50-70% and 80-90% confluency, respectively, after 24h incubation. Cells were transiently transfected using Lipofectamine LTX (Thermo Fisher Scientific) according to the manufacturer’s protocol. Transfection mixes were prepared with 1.25 µg pCAGGS.SARS-CoV-2_SΔ19_fpl_mNG2(11)_opt plasmid and 1.25 µg pCAGGS.BSD_fpl_mNG2(11) for HEK293T transfection; and 2.5 µg pcDNA3.1.mNG2(1-10) for A549.ACE2^+^ transfection. HEK293T cells were allowed to incubate for 24 h for efficient exogenous spike protein expression. At 6 h post transfection, transfected A549.ACE2^+^ cells were digested with 0.05% trypsin, washed, resuspended and counted on a Luna cell counter (Logos Biosystems), added to a 96-well plate at 2.2 × 10^4^ cells per well and incubated for 18 h. Transfected HEK293T cells were digested with 0.25% trypsin, washed, resuspended and then added to A549.ACE2^+^ cells at 2 × 10^4^ cells per well. Next, cells were imaged for 24 h using the IncuCyte® S3 Live-Cell Analysis System (Sartorius) at 20 min intervals. Image processing was performed using the IncuCyte® software.

### 2.10 Pseudovirus production

Production of VSV luciferase-based pseudovirus was done as follows. At first, HEK293T cells were seeded in a type I collagen-coated T75 flask in DMEM supplemented with 10% FBS at 3 × 10^6^ cells per flask. The following day, the HEK293T cells were transfected with 30 µg of expression plasmid encoding the SARS-CoV-2 Wuhan-Hu-1 S protein (pCAG3.1/SARS2-Sd19) or Delta variant S protein (pUNO1-SpikeV8; Invivogen) using FuGENE HD transfection reagent (Promega, Madison, WI, USA) in a 3:1 FuGENE HD:DNA ratio. Cell transfection was allowed for 24h at 37°C, 5% CO_2_. On day 3, serum-containing medium was replaced by serum-free DMEM and cells were inoculated with MOI 3 of VSVΔG*/Luc-G (Kerafast, Boston, MA, USA) for the production of luciferase-expressing pseudotypes. Incubation with virus-containing medium was allowed for 1h at 37°C, 5% CO_2_. Thereafter, cells were gently washed once with PBS and fresh DMEM/10% FBS with anti-VSV-G antibody (1:1000) was added for overnight pseudovirus production. Cell culture supernatants containing VSV pseudotyped with SARS-CoV-2 S protein (VSV-SARS2-Sd19/Luc) were collected at 24h post-infection. Finally, supernatants were centrifuged for 10 min at 1000 *g* to remove cell debris, and filtered once through a 0.45 µm pore size filter. Cleared supernatants containing SARS-CoV-2 S protein pseudotyped VSV particles were stored at -80°C.

SARS-CoV-2 VLP production for GFP read-out was carried out as follows. Briefly, HEK293T cells were seeded in a 165 cm^2^-dish 24 h before transfection. Upon transfection, the HIV backbone plasmid (pCAGGs Gag-Pol), a reporter plasmid (pQCXIP-GFP) and a spike-expressing plasmid (pCAGGS-SARS-CoV-2-spike) were co-transfected into HEK293T cells using Fugene HD transfection reagentia (Promega) according to the manufacturer’s guidelines. After 24h incubation (37°C, 5% CO_2_), culture supernatant was discarded and fresh DMEM supplemented with 8% inactivated FBS and 1 mM sodium butyrate was added. After another 24h incubation period at 37°C, 5% CO_2_, cell supernatant containing VLPs was collected and centrifuged at 1731 *g* for 10 min at 25°C. Then, supernatant was diluted (4 to 1 ratio *v/v*) with PEG-*it* solution (SBI, System Biosciences), vortexed and incubated continuously rotating overnight at 4°C. After 24h, the VLP solution was centrifuged for 30 min at 3000 *g* at 4°C and the resulting pellet was resuspended in one-tenth of the original supernatant volume with DMEM/10% FBS (heat-inactivated). SARS-CoV-2 VLPs were aliquoted and stored at -80°C.

### 2.11 Pseudovirus transduction assay

Pseudovirus transduction using (luciferase-based) SARS-CoV-2 spike pseudotyped VSV particles was performed as follows. A549.ACE2^+^.TMPRSS2^+^ target cells (Invivogen) were seeded in a white, clear-bottom 96-well plate at 1×10^4^ cells/well. After overnight incubation, compounds serially diluted (2X) in cell culture medium (DMEM/10% FBS) were added to the target cells. SARS-CoV-2 Wuhan-Hu-1 or Delta variant spike pseudotyped VSV (PV) particles were added to the target cells to reach a final infectious dose corresponding to the CCID_50_ of the virus stocks. Plates were then incubated at 37°C and 5% CO_2_ for 22h to allow infection. The following day, supernatant was removed from the target cells and Bright-Glo assay reagent (Promega) was added and, after a 5 min incubation period at RT, luminescence was detected on a GloMax Navigator microplate reader (Promega).

Transduction with (GFP-based) SARS-CoV-2 spike pseudotyped VLPs was done as follows. One day prior to start of experiment, A549.ACE2^+^ cells were first transfected with a plasmid expressing the TMPRSS2 gene. The next day, serial diluted compound was first incubated with VLPs for 30 min at 37°C, before this mixture was added to the A549.ACE2^+^-TMPRSS cells seeded in 96-well plates. Four days after pseudovirus transduction, immunofluorescent images were captured with a EVOS M5000 microscope (Invitrogen, Thermo Fisher Scientific) using the EVOS GFP led cube, employing a 4X UPlanSApo objective. After images were captured, pseudovirus-infected cells were counted using the default microscope software, selecting a target and background noise. Results are expressed as % inhibition compared to a virus control condition.

### 2.12 Flow cytometry

Intracellular nucleocapsid staining of infected cells was done as previously described (31). Briefly, infected cells were collected, washed in PBS and centrifuged in a cooled centrifuge (4°C) at 500 *g* for 5 min. After removal of the supernatant, cells were stained using a Fix/Perm kit (cat n° 554714, BD Biosciences). Cells were first fixed and permeabilized by the addition of 250 µL of BD Cytofix/Cytoperm buffer and incubated 20 min at 4°C. Samples were then washed twice with Perm/Wash buffer before the addition of the primary (anti-nucleocapsid) antibody (0.3 µg per sample). After a 30 min incubation at 4°C, samples were washed twice in BD Perm/Wash buffer, followed by a 30 min incubation at 4°C with the secondary (labeled) antibody, and washed again. Finally, samples were stored in PBS/2% PFA. Sample acquisition was done on a BD FACSCelesta flow cytometer (BD Biosciences) with BD FACSDiva v8.0.1 software. FACS data analysis (including cell debris and doublet exclusion) was done using FlowJo v10.1 (Tree Star).

Flow cytometric analysis of spike expression levels was performed as follows. Spike-transfected HEK293T cells were first digested using Trypsin-EDTA 0.05%, washed in phosphate buffered saline (PBS) with 2% FBS and resuspended at 3 × 10^6^ cells per ml. For each sample, 0.3 × 10^6^ cells were preincubated with spike-specific monoclonal antibodies in PBS/FBS 2% for 30 min at RT. The cells were washed once in PBS/FBS 2% before incubation with the appropriate species reactive and labeled secondary antibodies. Following incubation (30 min at RT), cells were washed twice, resuspended in PBS containing 1% paraformaldehyde and analysed on a FACSCelesta flow cytometer (BD Biosciences). Data analysis was done using FlowJo v10.1 software (Tree Star).

### 2.13 Surface plasmon resonance

SPR technology (Biacore T200, Cytiva) was used to determine the binding kinetics and affinity of UDA to the wild-type Wuhan-Hu-1 (2019-nCoV spike protein, cat n° MBS8574721, Mybiosource) and Omicron (COV2 spike protein S recombinant B.1.1529 Omicron, cat n° MBS553745, Mybiosource) SARS-CoV-2 spike protein, as well as to the Wuhan-Hu-1 SARS-CoV-2 receptor binding domain (RBD). For the binding study with UDA, RBD (2019-nCoV spike RBD, cat n° 40592-VNAH, SinoBiological) was immobilized on a CM5 sensor chip using standard amine coupling in 10 mM HEPES (pH 7.0) to a level of approximately 300 RU. For the additional binding study with the spike-binding antibodies, histidine-tagged RBD (2019-nCoV spike RBD his tag, cat n° 40592-V08H, SinoBiological) was used. Histidine-tagged proteins were capture-coupled an a nitrilotriacetic acid (NTA) sensor chip (Cytiva). Briefly, the NTA surface was first activated with 0.5 mM Ni^2+^ followed by a mixture of EDC/NHS to activate the carboxyl groups. Histidine-tagged proteins were diluted in HBS-P^+^ (10 mM HEPES, 150 mM NaCl, 0.05% surfactant P20; pH 7.4) and capture-coupled onto the surface to a level of 100-300 RU. Finally, the surface was deactivated using 1.0 M ethanolamine-HCl pH 8.5 and regenerated with 350 mM EDTA to remove any remaining unbound ligand. Interaction studies between UDA and spike/RBD were performed at 25°C in HBS-EP^+^ (10 mM HEPES, 150 mM NaCl, 3 mM EDTA, 0.05% surfactant P20; pH 7.4). Two-fold serial dilutions of UDA were injected at 30 µl/min using multiple cycle kinetics. 10 mM NaOH was used to regenerate the surface. Several buffer blanks were included for double referencing. The neutralizing R001 Ab and the non-neutralizing R007 Ab were used as positive controls for RBD binding. In addition, SPR technology was used to determine the inhibitory potential of UDA on the RBD/ACE2 binding. A biotin CAPture kit (Cytiva) was used to reversible capture biotinylated ACE2 (SinoBiological) in HBS-EP^+^ running buffer. The CAP sensor chip was first activated by injecting the Biotin CAPture reagent for 240 seconds (2 µl/min). Biotinylated ACE2 was captured onto the chip by injecting it for 180 seconds at a concentration of 5 µg/ml (10 µl/min). RBD (50 nM) alone or premixed with 1 µM UDA was injected for 120 sec (30 µL/min). RBD (50 nM) was also mixed with the spike neutralizing R001 Ab and the non-neutralizing R007 Ab at equimolar ratios. The surface was regenerated using the regeneration mix according to the manufacturer’s instruction. Several buffer blanks were included for double referencing.

Apparent binding kinetics (*K*_D_, *k*_a_, *k*_d_) were derived after fitting the experimental data to the 1:1 Langmuir binding model in the Biacore T200 Evaluation Software 3.1. The experiments were performed at least in duplicate.

### 2.14 Statistical analysis

Data were visualized as means ± standard deviation (SD) and were analyzed by making use of the GraphPad Prism 9.3.1 software.

## 3 Results

### 3.1 Antiviral activity of UDA against pseudotyped SARS-CoV-2

The plant lectin UDA was initially evaluated against pseudotyped SARS-CoV-2. First, lung epithelial A549 cells were stably transduced with a lentivector encoding the ACE2 receptor, to enhance their sensitivity to SARS-CoV-2, as described recently for the U87 cell line (30). As shown in **Supplementary Figure 1**, the resulting A549.ACE2^+^ cells expressed high and stable ACE2 levels as evidenced by the dense protein bands on the immunoblot. Next, these A549.ACE2^+^ cells were transiently transfected with a plasmid encoding the cellular protease TMPRSS2, which promotes viral entry through plasma membrane fusion (9).

**Figure 1.**
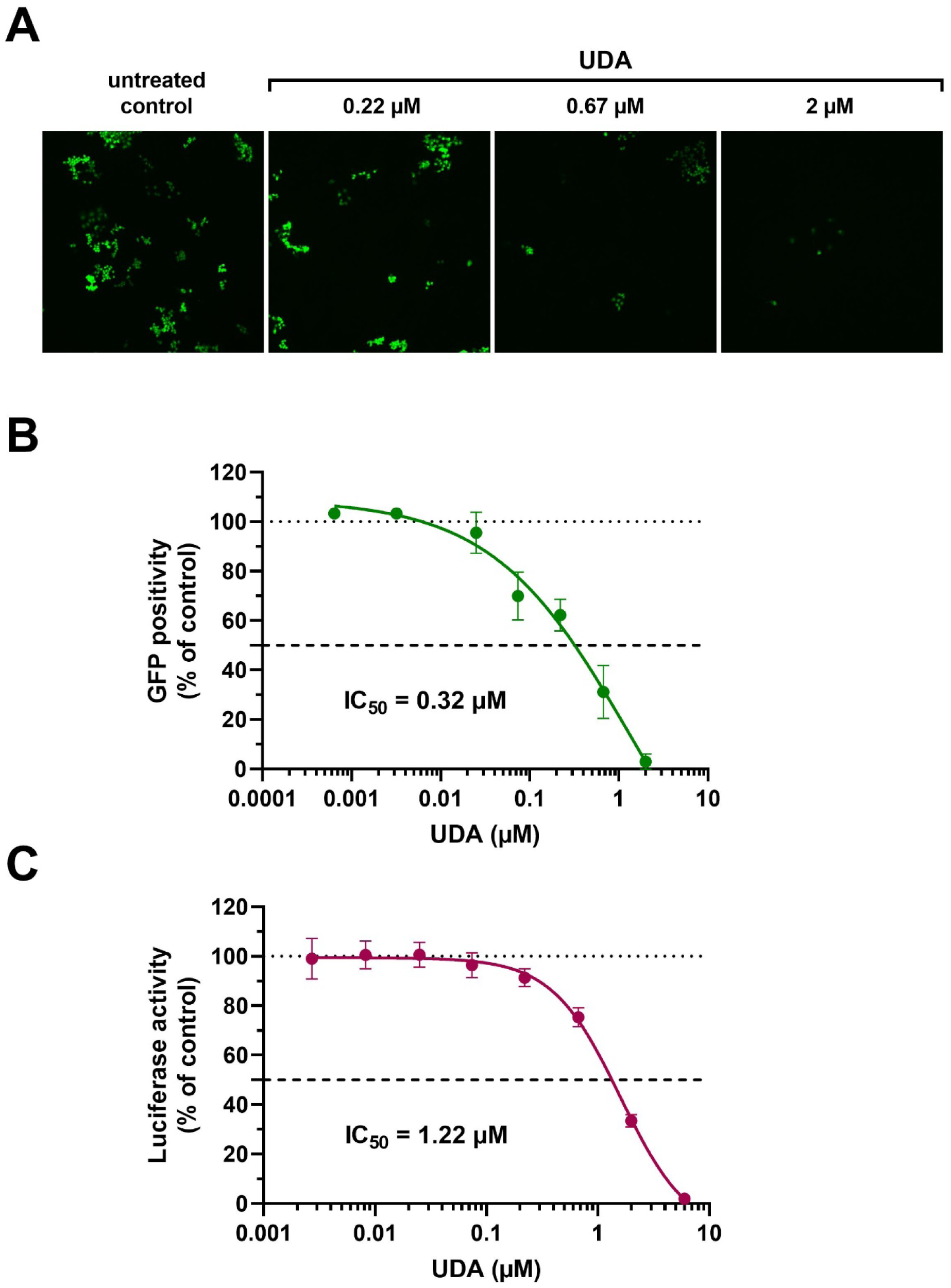
Antiviral activity of UDA against pseudotyped SARS-CoV-2. VLPs expressing the Wuhan-Hu-1 spike and a GFP or luciferase reporter was used to transduce cells in the absence (untreated control) or presence of UDA (as indicated). (**A**) A549.ACE2^+^ cells were transiently transfected with TMPRSS2 and subsequently exposed to pseudotyped SARS-CoV-2 expressing GFP. Panel shows representative images taken at 3 days post-transduction. (**B**) GFP-positive cells were quantified from images of *(A)*. Graph represents a concentration-response of UDA (mean ± SD from 3 independent experiments). (**C**) UDA was tested against luciferase-based pseudotyped SARS-CoV-2 in commercially available A549.ACE2^+^.TMPRSS2^+^ cells. Luciferase activity was measured at 22h after VLP transduction. Graph represents a concentration-response of UDA from 3 biological replicates in quadruple (mean ± SD; n=12).

The A549.ACE2^+^-TMPRSS2 cells were subsequently transduced with SARS-CoV-2 virus-like particles (VLPs) that carried the spike protein of the early Wuhan-Hu-1 strain and a GFP reporter. Interestingly, SARS-CoV-2 pseudovirus entry, evidenced by GFP expression in the transduced cells (**Figure 1A**), was profound and concentration-dependent inhibited by UDA, returning an IC_50_ value of 0.32 µM (**Figure 1B**). In addition, UDA activity was confirmed in a luciferase-based assay in A549.ACE2^+^.TMPRSS2^+^ cells (**Figure 1C**), showing a comparable, slightly higher IC_50_ value of 1.22 µM.

### 3.2 Broad spectrum antiviral activity of UDA against SARS-CoV-2 variants of concern

Next, the antiviral potency of UDA was evaluated against wild-type virus, as described recently (31). Briefly, Vero E6 cells were infected with clinical isolates of SARS-CoV-2 in the presence of UDA, and viral RNA was measured in the supernatant at day 3 post infection (p.i.). As shown in **Figure 2A** and **2B**, UDA inhibited SARS-CoV-2 replication in Vero E6 cells in a concentration-dependent manner, with IC_50_ values in the high nanomolar range (115-225 nM; **Table 1**). Importantly, UDA demonstrated antiviral activity against all tested variants, including Wuhan-Hu-1, 20A.EU2, and variants of concern (VOCs) Alpha (UK), Beta (South-African) and Gamma (Brazilian) (**Table 1**). At a concentration of 2 µM, UDA fully protects against SARS-CoV-2 infection, as confirmed by flow cytometric analysis of viral N protein expression in Vero E6 cells infected with the Gamma strain (**Figure 2C**).

**Figure 2.**
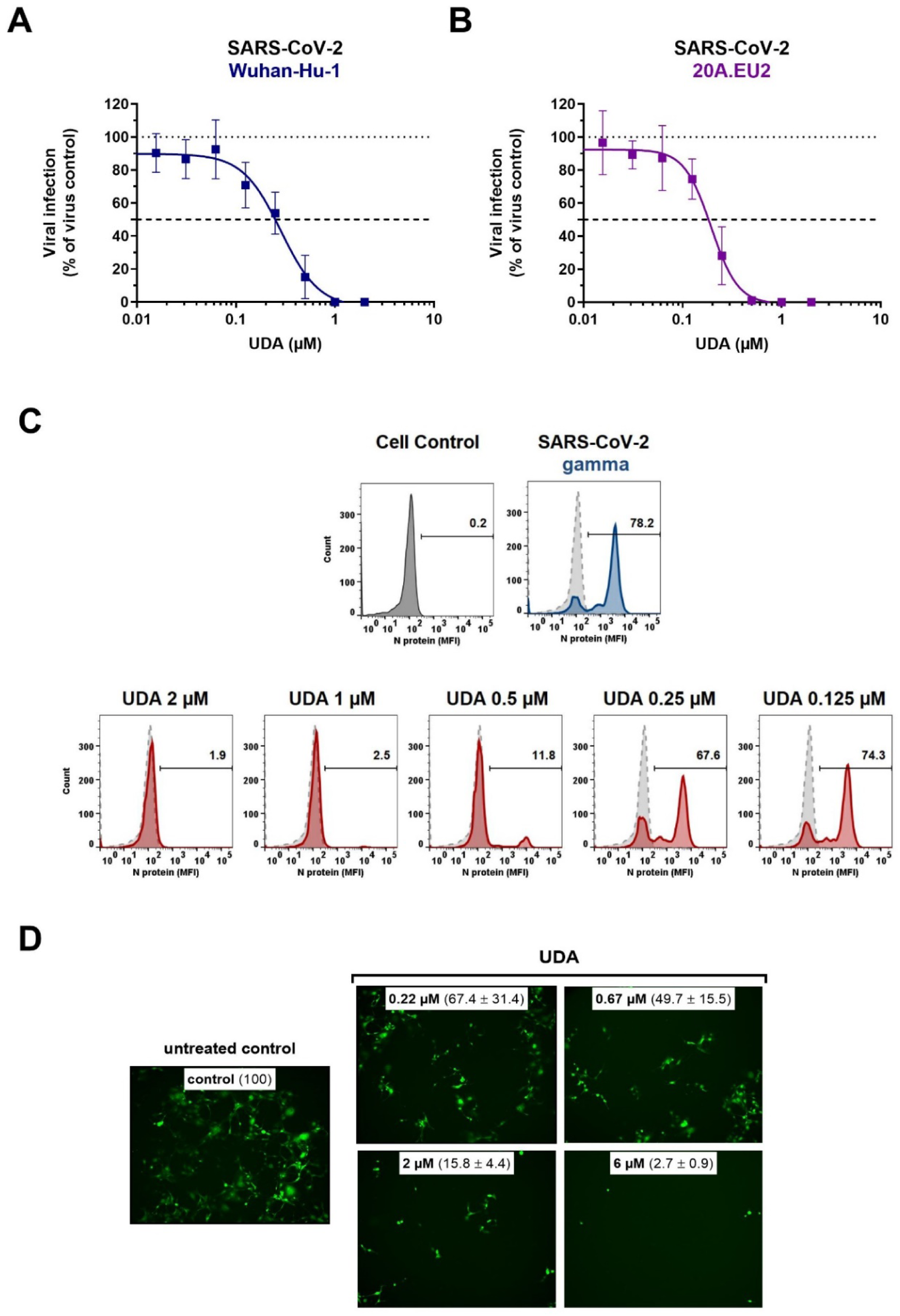
Antiviral activity of UDA against live SARS-CoV-2 virus in Vero E6 and U87.ACE2^+^ cells. Cells were exposed to clinical isolates of SARS-CoV-2, i.e., variants Wuhan-Hu-1 (A), 20A.EU2 (B), Gamma (C) or GFP-expressing Wuhan-Hu-1 (D) in the absence or presence of UDA. (**A** and **B**) SARS-CoV-2 replication was assessed by RT-qPCR analysis of the viral copy numbers of the N gene in the supernatant at day 3 post infection (p.i.). RT-qPCR data were used to calculate the % inhibition of viral replication and to plot a concentration-response curve for UDA. Graphs show data of 3 independent experiments with 2 technical replicates each (mean ± SD; n=6). (**C**) Cells were collected at 40h p.i. and stained intracellularly for the viral N protein. Histogram plots show mean fluorescence intensity (MFI) values of N expression in noninfected (Cell Control; grey), infected (Virus Control; blue) and UDA-treated infected (red) Vero E6 cells from a representative experiment. Single cell analysis was performed on 8,000 – 10,000 cells by flow cytometry. The numbers in each plot refer to the percentage of cells that stained positive for N (i.e., infected cells). The dashed grey histogram plot represents the background signal from the non-infected cell control. (**D**) Pictures, taken at 2 days post infection, show GFP expression in the infected U87.ACE2^+^ cells. Representative pictures from a biological replicate out of two are shown. The values between brackets refer to the percentage GFP^+^ area (relative to the virus control); mean ± SD (n=2).

**Table 1.** Antiviral activity of UDA against live SARS-CoV-2 virus in different cell lines.

| SARS-CoV-2 strain | UDA IC<sub>50</sub> (nM) |  |  |
| --- | --- | --- | --- |
|  | Vero E6<sup>a</sup> | U87.ACE2<sup>+b</sup> | A549.ACE2<sup>+</sup>-T2<sup>a</sup> |
| Wuhan-Hu-1 | 225 ± 71 | 984 ± 353 | 39.6 ± 6.5 |
| 20A.EU2 | 160 ± 33 | nd | 96.1 ± 28.0 |
| Alpha | 115 ± 69 | nd | nd |
| Beta | 118 ± 73 | nd | nd |
| Gamma | 171 ± 60 | nd | nd |
| Delta | nd | 1555 ± 106 | nd |
| Omicron | nd | 867 ± 731 | nd |

<sup>a</sup>Antiviral activity determined by RT-qPCR quantification of viral copy numbers of N gene in supernatant of infected cells. Mean ± SD; n=3.
<sup>b</sup>Antiviral activity determined by cell viability readout with MTS to quantify virus-induced cytopathic effect. Mean ± SD; n=3, except for Delta for which n=2.
IC<sub>50</sub>: the median inhibitory concentration 50%, or the concentration that inhibited SARS-CoV-2 infection by 50%; nd: not determined; T2: TMPRSS2.

In addition, we also tested UDA against the latest circulating Delta and Omicron variant. In order to get successful infection of the cells with Omicron SARS-CoV-2, we employed the glioblastoma cell line U87.ACE2^+^ which is highly permissive to SARS-CoV-2, as recently reported (30). As shown in **Figure 2D**, UDA preserved antiviral activity against SARS-CoV-2 in the U87.ACE2^+^ cells as evidenced by the reduced GFP expression in the cells exposed to a GFP-expressing Wuhan-Hu-1 variant. However, UDA was less potent against wild-type Wuhan-Hu-1 in the U87.ACE2^+^ cells (IC_50_ value of 984 nM; **Table 1**) as compared to the Vero E6 cells (IC_50_ value of 225 nM). Importantly, UDA also exerted antiviral activity against the Delta and Omicron VOCs (IC_50_ values of 1555 nM and 867 nM, respectively; **Table 1**), although with slightly reduced potency against Delta as compared to Wuhan-Hu-1, which is in line with the data obtained with pseudotyped Delta SARS-CoV-2 (**Supplementary Figure 2**). Nevertheless, these results demonstrate a broad-spectrum antiviral activity, including against the circulating and more infectious SARS-CoV-2 species.

As SARS-CoV-2 infection of Vero E6 and U87.ACE2^+^ cells mainly occurs via the endosomal entry route, we next evaluated the antiviral activity of UDA in our A549.ACE2^+^-TMPRSS2 cells, where the virus follows the cell surface entry route. Here, UDA also yielded full protection of virus infection at the highest concentration, as determined by flow cytometry (**Figure 3A**) and RT-qPCR (**Figure 3B**), with IC_50_ values of 40 and 96 nM for variants Wuhan-Hu-1 and 20A.EU2, respectively (**Table 1**).

**Figure 3.**
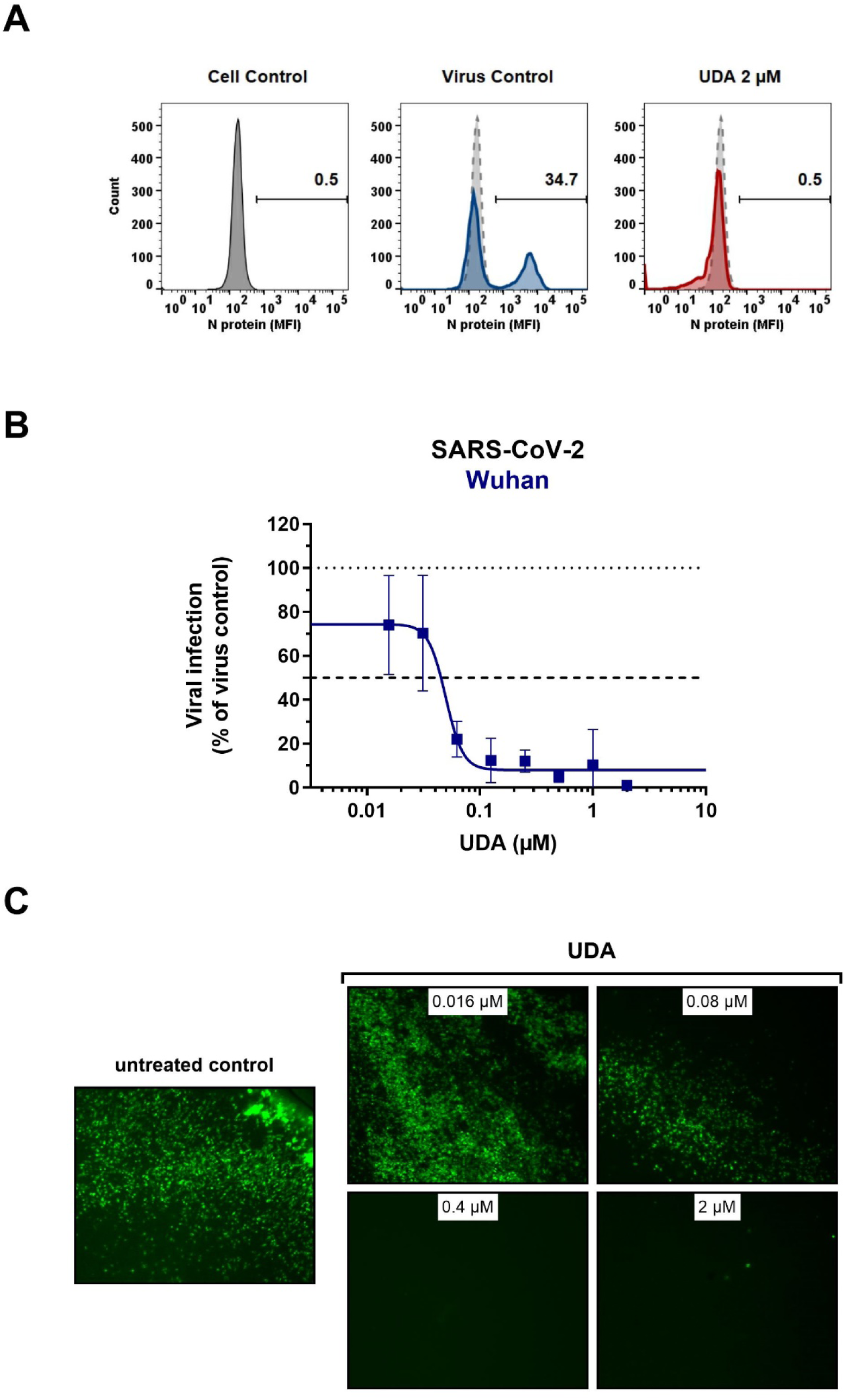
Antiviral activity of UDA against live SARS-CoV-2 virus in A549 cells and primary ALI cultures. (**A-B**) A549.ACE2^+^-TMPRSS2 cells were exposed to SARS-CoV-2 (WT Wuhan-Hu-1) in the absence or presence of UDA. (**A**) SARS-CoV-2 replication was assessed by flow cytometry. Cells infected with SAR-CoV-2 in the absence or presence of UDA (2 µM) were collected at 48h p.i. and stained intracellularly for the viral N protein. Histogram plots show mean fluorescence intensity (MFI) values of N expression in noninfected (Cell Control; grey), infected (Virus Control; blue) and UDA-treated infected (red) cells from a representative experiment. Single cell analysis was performed on 8,000 – 10,000 cells by flow cytometry. The numbers in each plot refer to the percentage of cells that stained positive for N (i.e., infected cells). The dashed grey histogram plot represents the background signal from the non-infected cell control. (**B**) SARS-CoV-2 replication was assessed by RT-qPCR analysis of the viral copy numbers of the N gene in the supernatant at day 3 post infection (p.i.). RT-qPCR data were used to calculate the % inhibition of viral replication and to plot a concentration-response curve for UDA. Graphs show data of 3 independent experiments with 2 technical replicates each (mean ± SD; n=6). (**C**) Human primary lower (SmallAir) airway epithelial ALI cultures were infected apically with a GFP-expressing SARS-CoV-2 variant (Wuhan-Hu-1) for 2h in the absence or presence of UDA, washed and exposed to air. Pictures, taken at 4 days post infection, show GFP expression in the infected cells. Representative pictures from a biological replicate out of two are shown.

To further address the antiviral potency of UDA, we evaluated its antiviral activity in differentiated cells in 3D like structures that are exposed to air, the so-called air-liquid-interface (ALI) cultures. In this experimental setting, the virus (in the absence or presence of compound) is only briefly (2h) added to the apical side of the cells, and viral replication is monitored by GFP expression or RT-qPCR at day 4 p.i. We used the human primary upper (MucilAir) and lower (SmallAir) airway epithelial cultures (from healthy donors), to create a more clinically relevant setting. Interestingly, even a short treatment of the virus with UDA at the time of infection strongly prevented the infection of ALI cultures of primary cells. As illustrated in **Figure 3C**, UDA yielded full protection of SARS-CoV-2 infection (GFP-expressing Wuhan-Hu-1 variant) in the SmallAir ALI cultures at 0.4 µM, as evidenced by the absence of GFP expression. Notably, RT-qPCR analysis of the apical washes revealed a stronger antiviral effect of UDA in SmallAir cultures infected with the 20A.EU2 variant (IC_50_ < 16 nM; more than 50% protection at the lowest tested UDA concentration of 16 nM), as compared to MucilAir cultures (IC_50_ = 272 ± 155 nM; n=2). To conclude, both viral entry routes can be efficiently blocked by UDA, which demonstrated a wide-spectrum antiviral activity against SARS-CoV-2 in different cell types.

### 3.3 Surface plasmon resonance analysis of UDA binding to SARS-CoV-2 spike protein

UDA is a carbohydrate binding agent, that has a preference for GlcNAc and high-mannose sugars on target glycoproteins (23, 24). As both the SARS-CoV-2 spike protein and ACE2 receptor are heavily glycosylated (37-40), UDA could possibly bind to both. Thus, we used surface plasmon resonance (SPR) to determine what domain of the viral S protein and/or cellular receptor is responsible for the antiviral activity of UDA. As expected, there was a clear concentration-dependent binding of UDA to monomeric spike protein of the early Wuhan-Hu-1 strain (**Figure 4A**), with a mean *K*D of 7 nM (**Supplementary Figure 3A)**, and also to the spike protein of the latest Omicron variant (mean *K*D of 11 nM; **Supplementary Figure 3A** and **3B**). However, the binding of UDA to the receptor binding domain (RBD) of the S protein was weaker as compared to full-length spike, with a fast off-rate (**Figure 4B**), indicative of a transient interaction of UDA to RBD (mean *K*D of 22 nM; **Supplementary Figure 3A)**. Furthermore, RBD with bound UDA could still bind to ACE2 (**Figure 4C**; slightly higher response of RBD + UDA compared to RBD alone). The same was seen for non-neutralizing control antibody R007, while spike-neutralizing antibody R001 completely blocked the binding of RBD to the ACE2 receptor (**Figure 4C** and **Supplementary Figure 3C**), in line with its antiviral activity against SARS-CoV-2 in Vero E6 cells, as recently described (31). In addition, UDA did not bind to the ACE2 receptor (**Figure 4C**). These data demonstrate that UDA is not acting as a direct receptor-attachment competitor, and its strongest interaction site is not located in the RBD.

**Figure 4.**
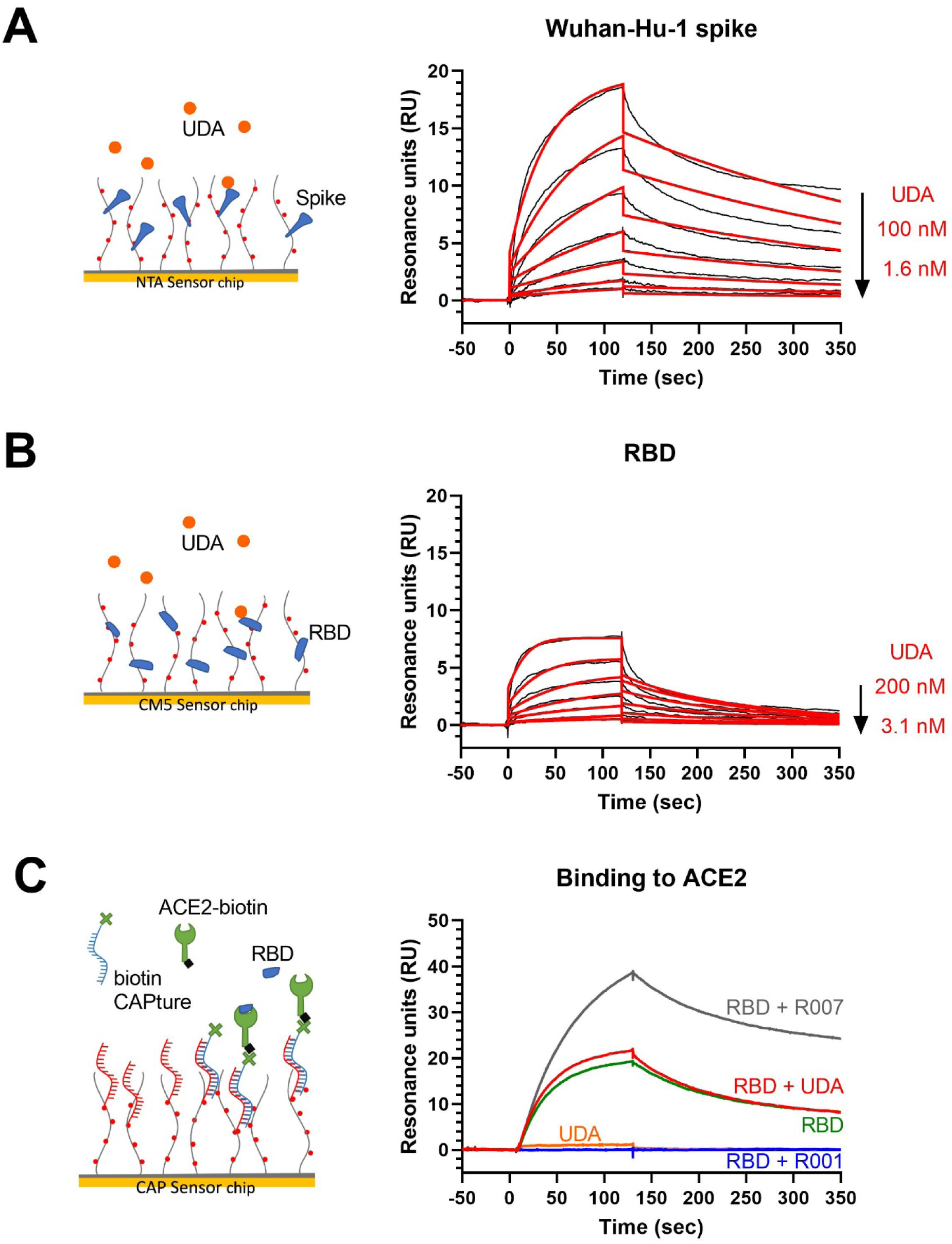
Surface plasmon resonance (SPR) analysis of UDA. (**A**) SPR sensorgram showing the binding kinetics for UDA and immobilized monomeric Wuhan-Hu-1 spike protein (1:2 dilutions of UDA, starting from 100 nM). Data are shown as black lines, and the best fit of the data to a 1:1 binding model is shown in red. (**B**) SPR sensorgram showing the binding kinetics for UDA and immobilized RBD of Wuhan-Hu-1 spike protein (1:2 dilutions of UDA, starting from 200 nM), with a fast off rate. Data are shown as black lines, and the best fit of the data to a 1:1 binding model is shown in red. (**C**) Biotinylated ACE2 was coupled as ligand to a CAP sensor chip. Graph shows sensorgrams for the binding of different analytes to ACE2. Green curve: RBD (50 nM) only; red curve: RBD (50 nM) + UDA (1 µM); grey curve: RBD (50 nM) + non-neutralising spike-binding antibody R007 (50 nM); blue curve: RBD (50 nM) + spike-neutralising antibody R001 (50 nM); orange curve: UDA (1 µM) only. Note that RBD in complex with R007 can still bind to ACE2 resulting in a stronger resonance signal induced by the large protein complex. See Supplementary Figure 3A for kinetics values.

### 3.4 UDA inhibits SARS-CoV-2 spike-mediated cell-cell fusion

To further investigate the specific molecular target of UDA in SARS-CoV-2 entry, we next tested the potential of UDA in preventing cell-cell fusion by means of a split neongreen molecular system (**Supplementary Figure 4A**). Here, one part of neongreen (i.e., the first 10 beta-sheets) is expressed in the cytosol of A549.ACE2^+^ acceptor cells, and the other part (i.e., the remaining 11^th^ beta-sheet) is co-expressed with spike protein in HEK293T donor cells. As shown in **Figure 5**, a profound cell-cell fusion occurred in the control condition with the generation of multinucleated giant cells, as evidenced by the abundant neongreen expression. Cell-cell fusion was already visible within a few hours after cell overlay (see also **Supplementary movie**). In the presence of 5 µM of UDA, only few neongreen-positive syncytia could be observed, and the syncytia remained small in size, indicative of limited cells that were involved in syncytium formation (**Figure 5**). The inhibitory effect of UDA on cell-cell fusion was concentration-dependent. As expected, control HEK293T cells transfected with only the 11^th^ beta-sheet (thus, not expressing the spike protein) were not capable to fuse with the complementary ACE2-positive cells (**Figure 5**).

**Figure 5.**
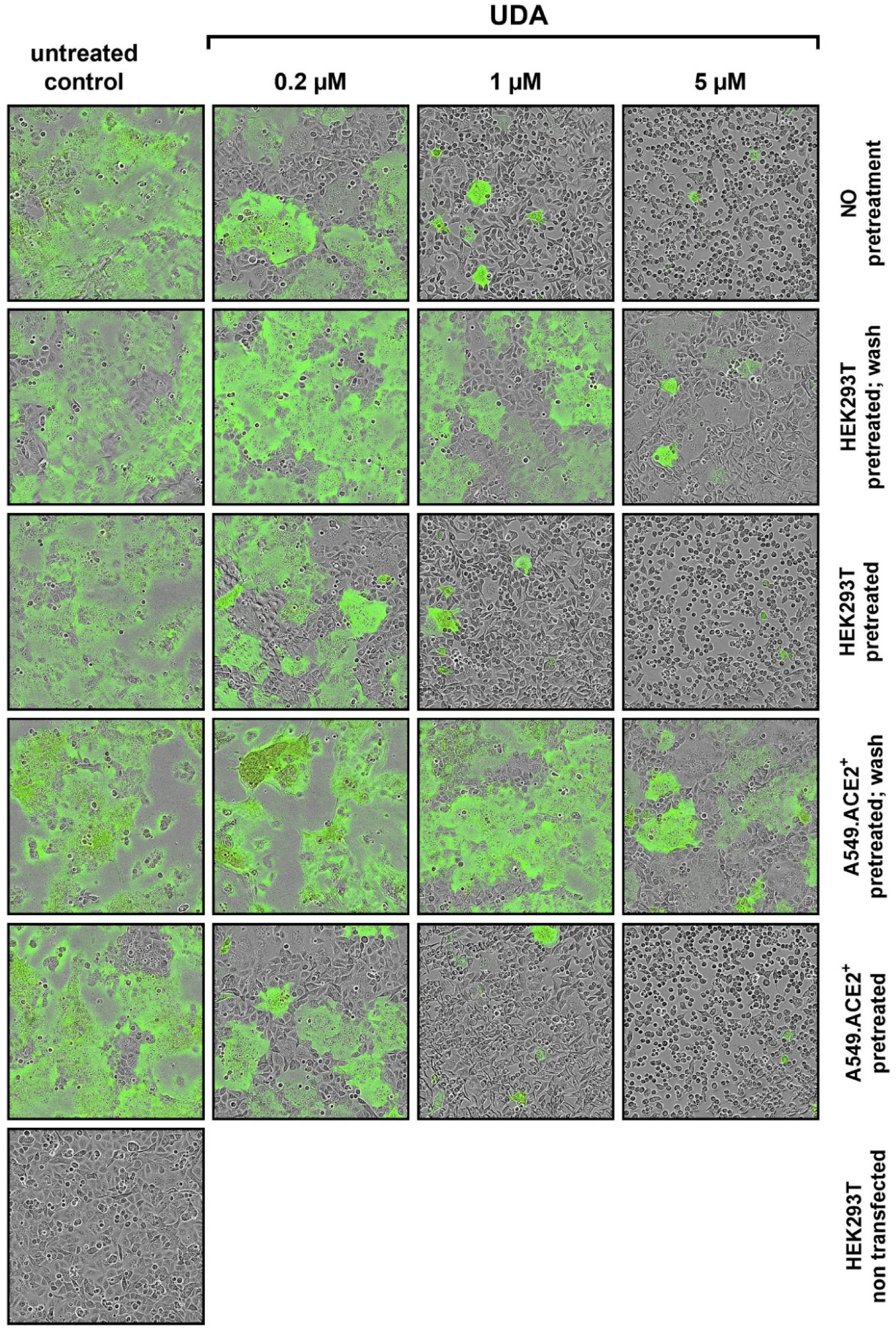
UDA prevents cell-cell fusion of A549.ACE2^+^ cells with spike-expressing HEK293T cells. A549.ACE2^+^ cells (transfected to express the first 10 betasheets of neongreen) were overlayed with HEK293T cells co-transfected with a plasmid encoding the SARS-CoV-2 spike protein and a plasmid encoding the 11^th^ betasheet of neongreen. Overlay was done in the absence (untreated control) or presence of UDA. Representative pictures of cell-cell fusion were taken 12h after the co-cultivation of both cell types. Top row (‘no pretreatment’): compound was added at the same moment as cell overlay; Second row (‘HEK293T pretreated; wash’): HEK293T were pretreated with UDA for 30 min; extensively washed and added to the A549 cells without compound; Third row (‘HEK293T pretreated’): HEK293T were pretreated with UDA for 30 min and added to the A549 cells with compound; Fourth row (‘A549.ACE2^+^ pretreated; wash’): A549 cells were pretreated with UDA for 30 min; extensively washed before the HEK293T cells were added without compound; Fifth row (‘A549.ACE2^+^ pretreated’): A549 cells were pretreated with UDA for 30 min before HEK293T cells were added without removal of compound; Bottom row (‘HEK293T non-transfected’): as a negative control HEK293T cells were transfected with only the 11^th^ betasheet of neongreen (without spike protein) and were added to the A549.ACE2^+^ cells (transfected with the first 10 betasheets of neongreen). For each condition, 2 replicate wells were analysed and in each well 4 different areas of the cell culture were monitored using an Incucyte live-cell analysis instrument. Representative pictures are shown. Neongreen expression analysis of the pictures is summarized in Supplementary Figure S3.

Whereas treatment of the HEK293T cells with UDA before the overlay on A549.ACE2^+^ cells prevented syncytia formation, pretreatment of the spike-transfected HEK293T cells with UDA (and removal of unbound UDA) did not inhibit cell-cell fusion to the same extent (**Figure 5**). The 5 µM UDA treatment did reduce the syncytia, as evidenced by the limited expression of neongreen over time (**Supplementary Figure 4B**), whereas 1 µM of UDA failed to prevent cell-cell fusion. Pretreatment of the spike-transfected HEK293T cells with UDA (without lectin wash-out) before the overlay on the A549.ACE2^+^ cells had no additional effect on fusion inhibition (**Supplementary Figure 4B**; compare 0.2 µM UDA samples). These results indicate that the presence of UDA is required during the fusion process to exert its inhibitory effect. Also, pretreatment of the A549.ACE2^+^ acceptor cell monolayer with 5 µM of UDA (and removal of unbound UDA) did not inhibit cell-cell fusion, as only a small reduction in neongreen signal was observed (**Figure 5** and **Supplementary Figure 4B**). Hence, putative binding of UDA to the cellular receptors (and/or cell surface of the target cells) is not sufficient to prevent membrane fusion elicited by the SARS-CoV-2 spike protein.

### 3.5 Analysis of UDA interaction with glycan mutants of SARS-CoV-2 spike protein

Finally, we wanted to investigate which carbohydrates on the spike protein are involved in UDA binding. Given that a strong effect of UDA was seen on spike-mediated cell-cell fusion, we primarily analysed the contribution of the glycans on the S2 subunit of the spike protein. To accelerate our analysis, we generated spike mutants that contained two or three deletions of adjacent glycosylation sites (see scheme in **Figure 6**). We started with the construction and analysis of the following three mutants: N1074Q + N1098Q; N1134Q + N1158Q; and N1173Q + N1194Q. These mutant spike proteins were subsequently used to generate VLPs for transduction of A549.ACE2^+^-TMPRSS2 cells. As summarized in **Table 2**, UDA kept full activity against these mutant VLPs, suggesting that antiviral activity of UDA is not related to interaction with a single glycan in the C-terminal domain of the SARS-CoV-2 spike S2 subunit (**Figure 6**). Deletion of the three glycosylation sites in the N-terminal domain of the S2 subunit (i.e., mutant N709Q + N717Q + N801Q) resulted in low cell surface expression of the mutant spike protein (**Supplementary Figure 5**), and consequently, in unsuccessful production of pseudotyped virus. However, mutation N709Q in combination with deletion of the N234 glycosylation site (located in S1, and the only site within the spike that carries exclusively oligo-mannose glycans with up to 9 mannose residues (41)), resulted in comparable or even enhanced spike expression as compared to WT, depending on the specific anti-S antibody used (**Supplementary Figure 5**). As listed in **Table 2**, UDA kept full activity against this N234Q + N709Q mutant VLP. Finally, the N657 glycosylation site (in S1) was targeted, given that of the remaining glycosylation sites this glycan is positioned most closely to the stem of the S2 subunit (**Figure 6**). Also, for this N657Q mutant a clear antiviral effect on pseudovirus transduction was observed (**Table 2**). Thus, removal of the selected N-glycosylation sites of the spike protein had little impact on the antiviral effect of UDA.

**Figure 6.**
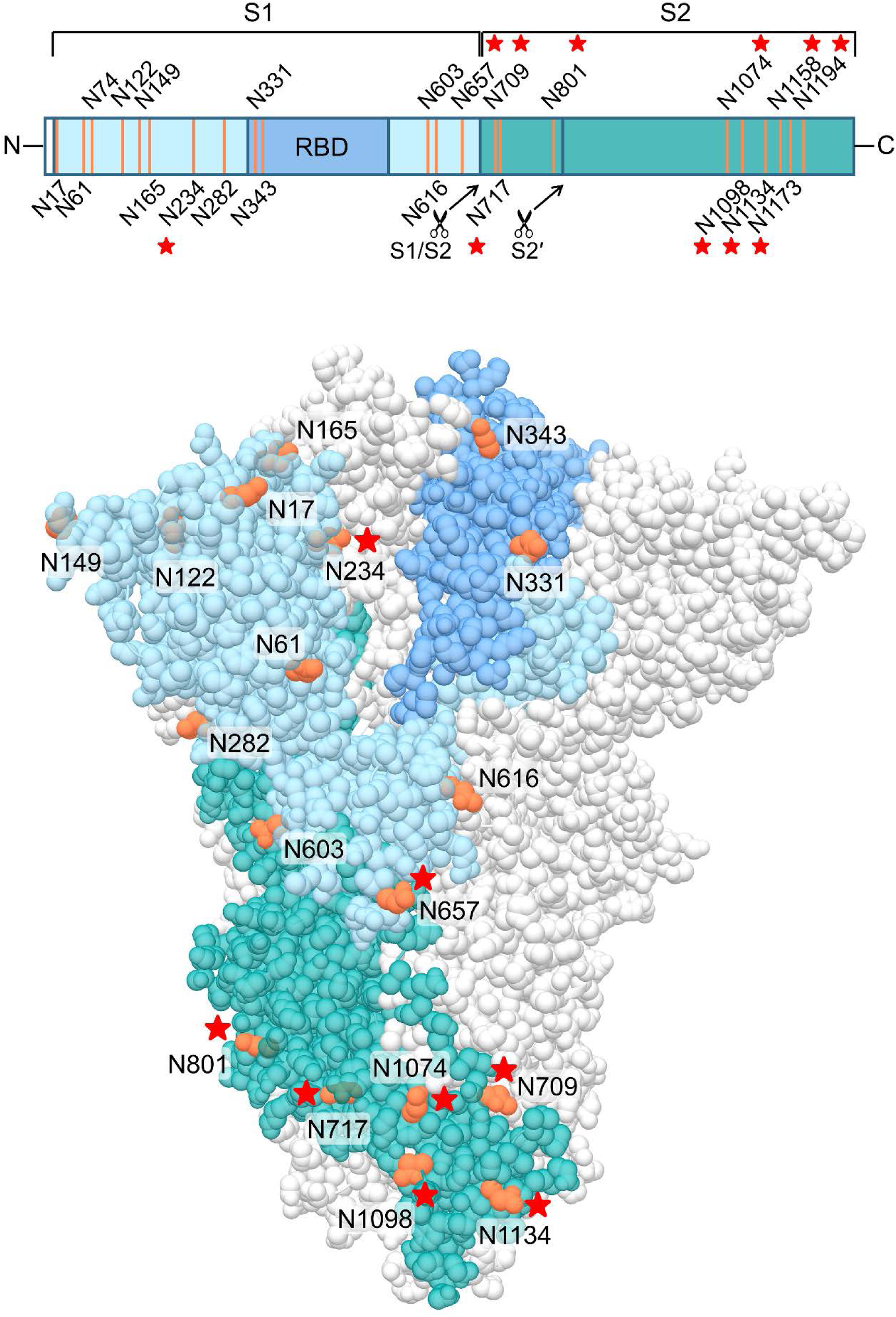
Structure of the SARS-CoV-2 spike trimer, based on PDB 6ZGE (56). One monomer is coloured, in blue (S1 subunit; with RBD in darker blue) and dark cyan (S2 subunit), with the N-glycosylation sites in orange. The stars indicate the N-glycosylation sites which were deleted in this study. Four N-glycosylation sites (i.e., N74, N1158, N1173 and N1194) were unresolved in the cryoEM structure. Image created with UCSF Chimera (57).

**Table 2.** Antiviral activity of UDA against glycosylation mutants of pseudotyped SARS-CoV-2.

| SARS-CoV-2 PV mutant | UDA IC<sub>50</sub> (nM)<sup>a</sup> |
| --- | --- |
| WT | 330 ± 125 |
| N1074Q; N1098Q | 157 ± 65 |
| N1134Q; N1158Q | 220 ± 115 |
| N1173Q; N1194Q | 146 ± 27 |
| N709Q; N717Q; N801Q | nd<sup>b</sup> |
| N234Q; N709Q | 390 ± 203 |
| N657Q | 435 ± 216 |

<sup>a</sup>Antiviral activity determined by GFP quantification of PV-transduced A549.ACE2<sup>+</sup>- TMPRSS2 cells. A549.ACE2<sup>+</sup> cells were transiently transfected with TMPRSS2 and subsequently exposed to pseudotyped SARS-CoV-2 expressing GFP. Mean ± SD; n=3, except for WT for which n=4 and S501 for which n=2.
<sup>b</sup>Mutant could not be analysed because of very low expression of spike protein.
nd: not determined

## 4 Discussion

Despite valuable progression in treatment and prevention of severe COVID-19, the persistent spread and rapid evolution of SARS-CoV-2 continue to give rise to new VOCs. Furthermore, genetic recombination of SARS-CoV-2 variants during co-infection could potentially further increase virulence, transmissibility and morbidity, especially in higher-risk individuals and immunocompromised patients (42, 43). In addition, the latest circulating SARS-CoV-2 VOCs, i.e., Delta and Omicron, have shown increasing resistance to SARS-CoV-2 RBD-specific neutralizing antibodies and current vaccines (44-46). Therefore, there is an urgent need for antiviral agents with potent anti-coronavirus activity and broad applicability. In this study, we evaluated the antiviral potential of UDA against SARS-CoV-2. From the data obtained with pseudotyped virus and live virus in different cell culture systems, we can conclude that UDA consistently inhibits entry of the virus into target cells. Our observation that UDA maintained antiviral activity among different SARS-CoV-2 VOCs suggests that UDA should be considered as an antiviral with an interesting pan-character that could serve as a valuable weapon in the combat against new emerging SARS-CoV-2 variants.

We could clearly demonstrate a profound inhibitory effect of UDA on SARS-CoV-2 spike-mediated fusion, as visualized with real-time microscopy. Targeting the first step in the viral life cycle, i.e., the engagement of host cell receptors and subsequent viral uptake, is an appealing antiviral strategy for several reasons. First, some entry mechanisms are widely conserved, even over different virus families. Second, entry inhibitors are not required to enter the cell because they interact with either a viral or a cell surface factor. This improves target accessibility and loosens restrictions in structural and chemical requirements, thereby allowing peptides and antibodies to be considered as drug candidates as well. Especially in the context of respiratory infections, nasal sprays can then be considered as an additional treatment option. Third, as blocking viral entry can prevent triggering the inflammatory cascade and avoid severe damage caused by the virus during a later stage in its life cycle, an entry inhibitor might improve disease outcome. Finally, entry inhibitors can potentially be used as both therapeutic and prophylactic drugs. The latter is especially interesting for healthcare workers and people traveling to endemic countries.

Previous work by Keyaerts *et al*. (29) already demonstrated a strong antiviral activity of UDA against SARS-CoV, most probably by hindering viral attachment. Plant lectins have also been shown to not only interfere with virus attachment for HIV (47), but also block virus-cell fusion for both HIV and influenza (25, 26, 48). Given that the basic principle for membrane fusion (e.g., heptad repeat domains and fusion peptide) in the fusion protein is conserved among different enveloped viruses, one can speculate that lectins, such as UDA, by interacting with glycans on the fusion protein might generally hamper the flexibility of the fusion protein to execute the fusion process. Our cell-cell fusion experiments clearly indicated that a saturating amount of UDA is required during the dynamic fusion process in order to evoke a (nearly) complete inhibitory effect. We observed that removal of unbound UDA before the initiation of cell-cell fusion resulted in a significant drop in the inhibition potential of UDA. This can either be because of a transient interaction of UDA with the S protein, which is not lasting long enough to prevent further fusion steps, or because UDA is acting at a specific step post receptor attachment by S, when a complete conformational change in the spike protein is taking place to execute the final steps in membrane fusion (e.g., detachment of S1 from S2, and the insertion of the fusion peptide in the host cell membrane with subsequent formation of the 6-helix bundle).

Via molecular docking it has been proposed that UDA specifically interacts with N-linked glycans on the RBD (49). However, our SPR data indicate that UDA is not directly interfering with binding of the RBD to ACE2, arguing for a post-attachment effect of UDA. Nevertheless, we cannot fully exclude an impact on spike attachment to ACE2, as in native trimeric spikes the ACE2 receptor-binding site is only exposed when the RBD is in the “up” conformation (50). Previous studies demonstrated that spike glycans, linked to N165, N234 (located outside the RBD) and N343 (located in the RBD), can modulate the RBD conformation. Removal of these glycosylation sites leads to a significant reduction of ACE2 binding, as the RBD will undergo a conformational shift towards the “down” state (51, 52). Thus, if UDA would target one of these glycans, which are located in or adjacent to the RBD, this could potentially alter the RBD conformation and therefore reduce the attachment efficiency of the virus to host cells. While substitutions N234Q and N657Q did not alter UDA activity, we did not assess the role of other S1 glycans in UDA binding yet. Alternatively, UDA could also interfere with viral entry by blocking binding to auxiliary receptors or cofactors, or by hampering protease cleavage at the S2′ site. More detailed analysis of the interaction of UDA on SARS-CoV-2 spike is needed to further elucidate its specific mode of action.

Containing 22 *N*-linked glycosylations on its surface, either complex type or oligomannose type glycans (40, 53, 54), SARS-CoV-2 spike presents multiple potential target sites for UDA interaction. Also, as UDA is one of the smallest plant lectins reported (21), it is not unlikely that multiple UDA molecules may simultaneously interact with the spike protein. Our initial glycosylation scan of the S2 subunit clearly shows that UDA activity is not related to a single *N*-glycosylation in the S2 subunit, as the removal of up to 3 glycosylation sites in S2 does not impact the antiviral activity of UDA. However, it is highly plausible that even more glycosylation sites on the spike protein need to be deleted before significant resistance against UDA could occur, as has been reported for HIV (55). Such mutant virus strains with a depleted glycan shield would become increasingly vulnerable to neutralising antibodies and the cellular immune system. In addition, loss of glycosylation has an impact on protein stability and functionality, and may render these escape mutants less infectious. This would suggest a high resistance barrier for UDA.

Taken together, our results demonstrate that UDA is a highly promising candidate for development as a potent and broadly acting antiviral agent against current and future SARS-CoV-2 variants.

## 6 Author Contributions

K.V., E.V., J.S. and S.N. conceived experiments; E.V., T.D., B.P., J.S., A.C. and S.N. performed experiments; K.V., E.V., J.S., S.N. and A.S. wrote the manuscript; D.S. secured funding; P.M. and D.S. provided reagents; E.V.D., P.M. and A.S. provided expertise and feedback.

## 8 Acknowledgments

We thank Geert Schoofs and Eef Meyen for their excellent technical assistance. Images were created with UCSF Chimera, developed by the Resource for Biocomputing, Visualization, and Informatics at the University of California, San Francisco, with support from NIH P41-GM103311; Graphpad Prism 9.3.1 (GraphPad Software, San Diego, California USA) and BioRender.com.

## 9 Supplementary Material

Supplementary Figures 1 – 5 and Supplementary Movie.

