## Supplementary figures for "Carbohydrate-Binding Protein from Stinging Nettle as Fusion Inhibitor for SARS-CoV-2 Variants of Concern"

*Supplementary Material*

### 1 Supplementary Figures

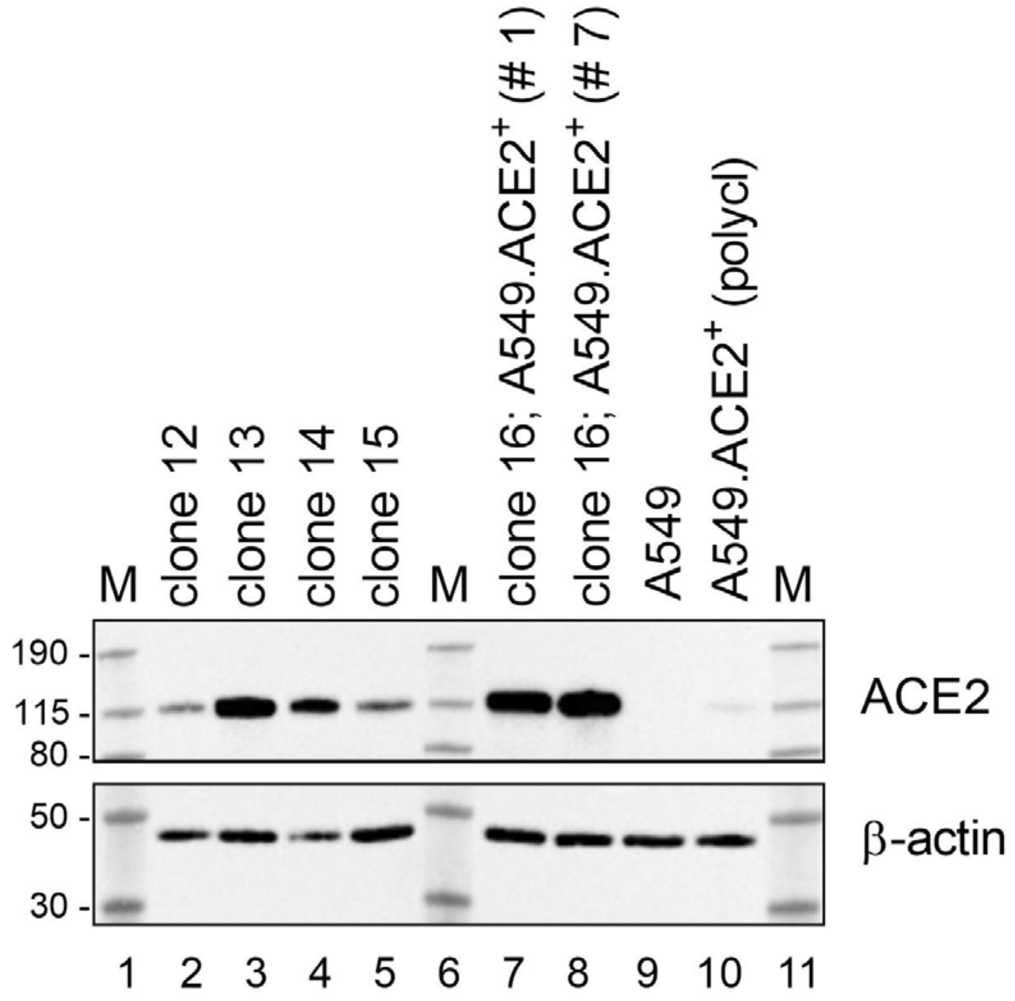


**Supplementary Figure 1.** Stable expression of ACE2 in transduced A549.ACE2$^{+}$ cells. Different clones of A549 cells were analysed for ACE2 expression by immunoblotting, with β-actin as loading control. Clone 16 was selected for this study. The stably ACE2 transduced A549 cells (clone 16) were tested at an early (# 1) and later (# 7) passage of the cells (lanes 7 and 8, respectively). The parental A549 cells did not express detectable levels of ACE2 (lane 9). The polyclonal mixture (polycl) of ACE2-transduced A549 cells is also included (lane 10), showing some expression of ACE2. M; molecular marker in kDa.

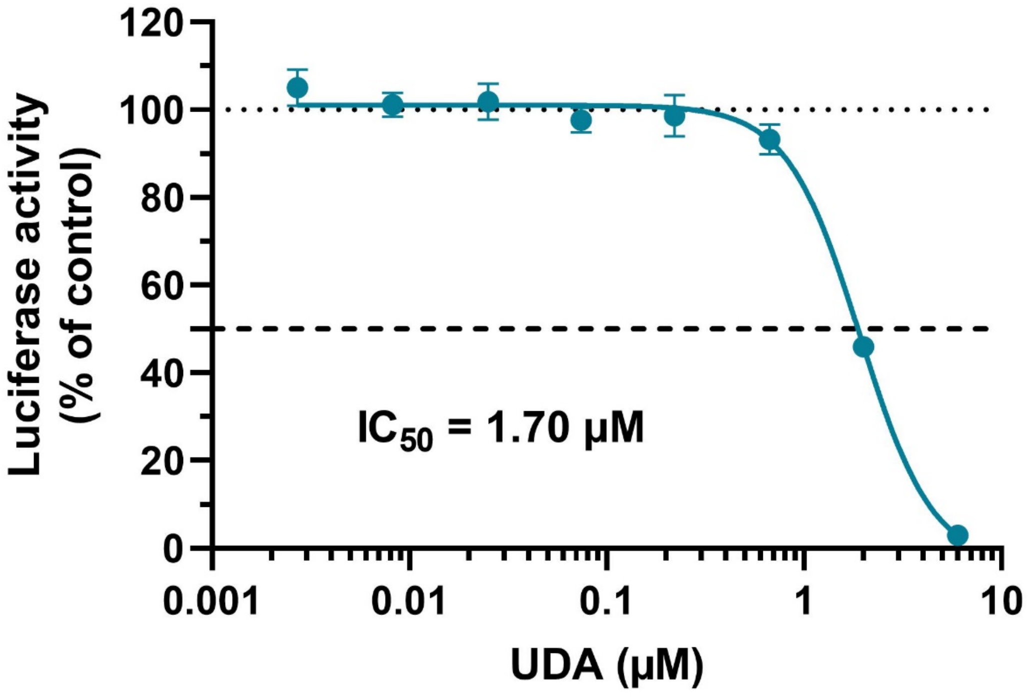


**Supplementary Figure 2.** Antiviral activity of UDA against pseudotyped Delta SARS-CoV-2. UDA was tested against luciferase-based pseudotyped SARS-CoV-2 (expressing the spike from the Delta VOC) in commercially available A549.ACE2$^{+}$.TMPRSS2$^{+}$ cells. At 22h after VLP transduction, luciferase activity was measured. Graph represents a concentration-response of UDA from 2 biological replicates in quadruple (mean ± SD; n=8).

**A**

| Ligand/Analyte | $k_a$ (x $10^5$ $M^{-1}s^{-1}$) | $k_d$ (x $10^{-5}$ $s^{-1}$) | $K_D$ (nM) |
|---|---|---|---|
| Wuhan-Hu-1 spike/UDA (n=8) | 3.37 ± 0.22 | 240.43 ± 13.03 | 7.37 ± 0.92 |
| Omicron spike/UDA (n=4) | 3.49 ± 0.20 | 364.55 ± 9.25 | 10.58 ± 0.82 |
| Wuhan-Hu-1 RBD/UDA (n=2) | 3.10 ± 0.60 | 709.50 ± 221.50 | 22.35 ± 2.85 |
| Wuhan-Hu-1 RBD/R001 (n=3) | 19.70 ± 0.87 | 4.17 ± 0.17 | 0.020 ± 0.001 |
| Wuhan-Hu-1 RBD/R007 (n=6) | 2.93 ± 0.37 | 16.80 ± 0.89 | 0.66 ± 0.15 |
| Wuhan-Hu-1 RBD/R007 (n=2)* | 4.00 ± 0.77 | 15.25 ± 1.35 | 0.39 ± 0.04 |

**B**

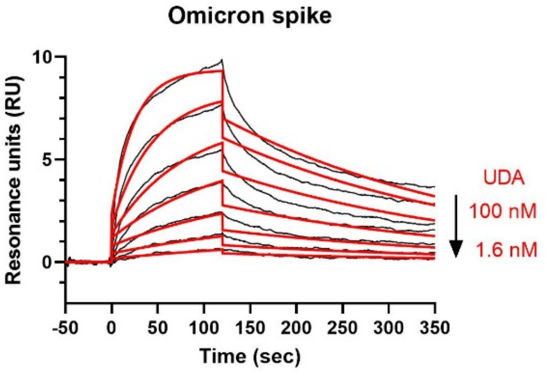


**C**

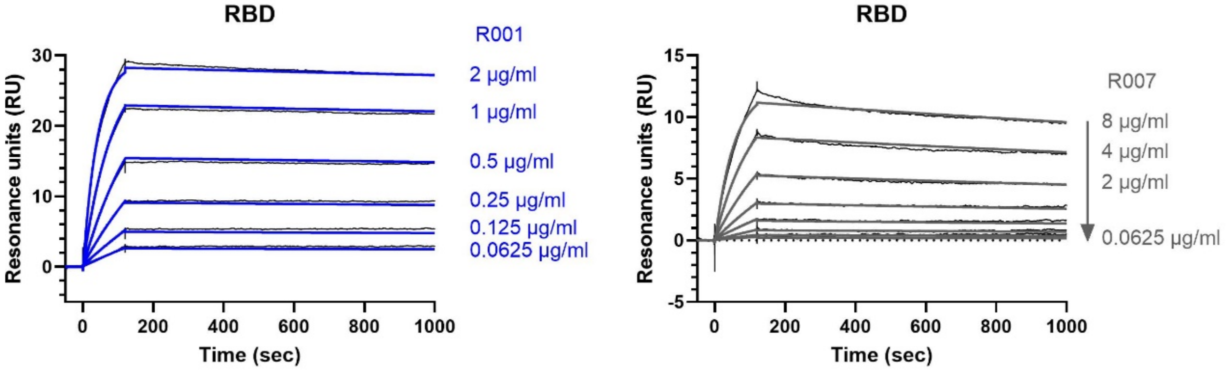


**Supplementary Figure 3.** Surface plasmon resonance (SPR) analysis of UDA and spike-binding antibodies R001 and R007. (**A**) Summary of the kinetics parameters from different repeat SPR experiments performed in this study. Given are the association rate constant ($k_a$), the dissociation rate constant ($k_d$), and the dissociation equilibrium constant ($K_D$). Values are mean ± SEM. (*) For the interaction of R007 to RBD, two different sensor chips were used, an NTA chip (n=8) or an CM5 chip (n=2). (**B**) SPR sensorgram showing the binding kinetics for UDA and immobilized monomeric Omicron spike protein (1:2 dilutions of UDA, starting from 100 nM). Data are shown as black lines, and the best fit of the data to a 1:1 binding model is shown in red. (**C**) SPR sensorgrams showing the binding kinetics for spike-binding antibodies and immobilized RBD of Wuhan-Hu-1 spike. Left panel shows sensorgram for the spike-neutralising antibody R001 and right panel that of the non-neutralising spike-binding antibody R007. Data are shown as black lines, and the best fit of the data to a 1:1 binding model is shown in blue or grey, respectively. SPR: surface plasmon resonance; RBD: receptor binding domain; UDA: *Urtica dioica* agglutinin; NTA: nitrilotriacetic acid.

**A**

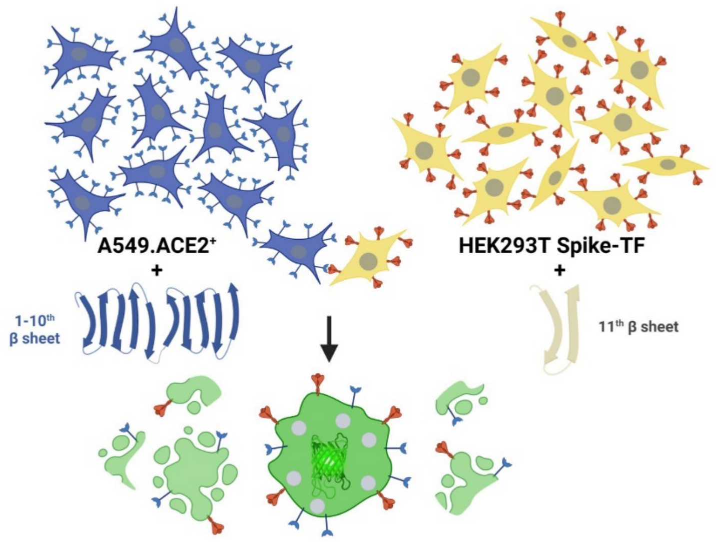


**B**

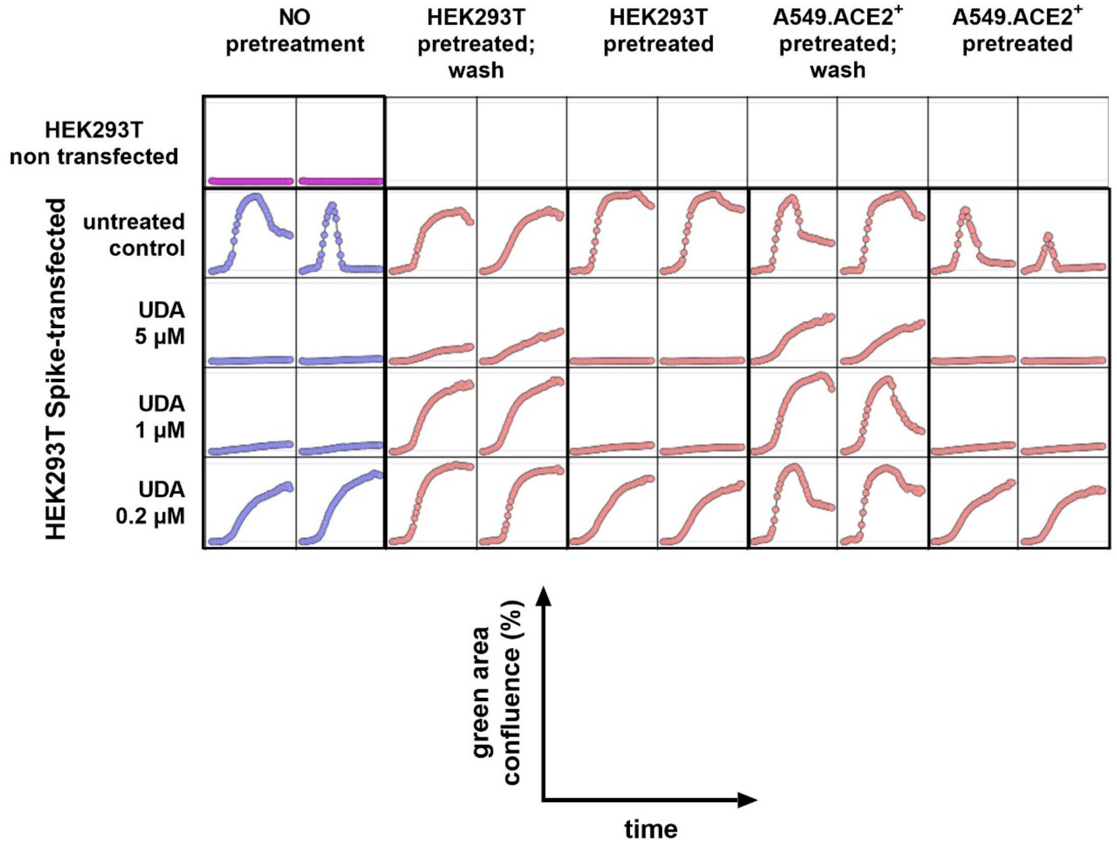


**Supplementary Figure 4.** UDA prevents cell-cell fusion of A549.ACE2$^{+}$ cells with spike-expressing HEK293T cells. (**A**) A549.ACE2$^{+}$ cells (transfected to express the first 10 betasheets of neongreen) were overlayed with HEK293T cells co-transfected with a plasmid encoding the SARS-CoV-2 spike protein and a plasmid encoding the 11$^{th}$ betasheet of neongreen. Only cell-cell fusion of an A549 cell with a HEK293T cell will result in the assembly of a functional neongreen protein and give a green fluorescence signal. (**B**) Samples from Figure 5 were analysed for neongreen expression (for specification of the samples, see legend to Figure 5). Each condition was tested in 2 replicate wells (side-by-side columns), and in each well 4 different areas of the cell culture were monitored using an Incucyte live-cell analysis instrument. The increase in neongreen expression over time (17 hours) is plotted as percentage of green area confluence. Each individual plot shows the average signal of the 4 different areas of the cell culture (mean; n=4). Note that because of lysis of the syncytia, the fluorescent protein is diluted in the culture medium, resulting in a drop in the neongreen signal at later time points.

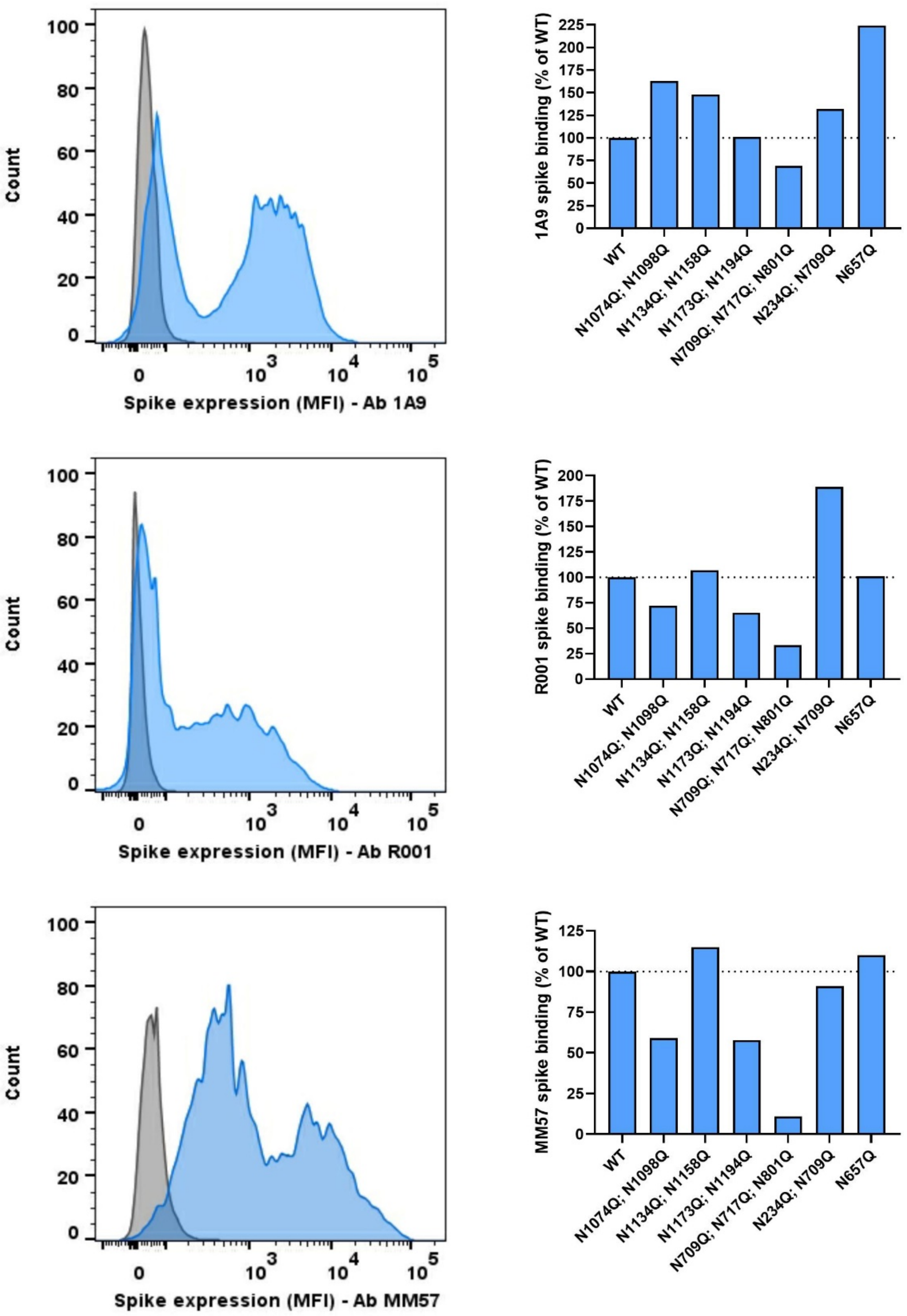


**Supplementary Figure 5.** Cell surface spike expression of different N-glycosylation mutants. HEK293T cells were transfected with a plasmid encoding the wild-type (WT) SARS-CoV-2 spike protein or an N-glycosylation deletion mutant. At 24h post transfection, cells were collected and stained with three different anti-spike antibodies (as indicated) to determine the cell surface expression of S.

Histogram plots on the left show the mean fluorescence intensity (MFI) of S protein expression for non-transfected (grey) and WT S-transfected (blue) HEK293T cells. Flow cytometric data were collected from approximately 6,000 analyzed cells. The bar graphs on the right show S expression for the different spike mutants as determined by staining with the corresponding anti-spike antibody (as indicated). Bars represent the MFI relative to the WT control.

### 2 Supplementary Movie

**Supplementary movie.** A549.$ACE2^+$ cells (transfected to express the first 10 betasheets of neongreen) were overlayed with HEK293T cells co-transfected with a plasmid encoding the SARS-CoV-2 spike protein and a plasmid encoding the $11^{th}$ betasheet of neongreen. Overlay was done in the absence (untreated control; left) or presence of UDA (1 µM; right). Fusion events were visualized using the IncuCyte® S3 Live-Cell Analysis System (Sartorius). Phase contrast and GFP images were taken using a 20x objective lens at 20 minute intervals for a 24 hours period. Image processing was performed using the IncuCyte software.
